# A high lipid diet leads to greater pathology and lower tolerance during infection

**DOI:** 10.1101/2024.08.02.606372

**Authors:** Weston G. Perrine, Erin L. Sauer, Ashley C. Love, Ashley Morris, Johnathon Novotny, Sarah E. DuRant

**Author notes:** Corresponding Author: Sarah E DuRant.

## Abstract

Despite the clear effect of food resources on wildlife disease dynamics, few studies have investigated how shifts in diet, specifically macronutrient content, shape vertebrate host responses to infection. While each individual macronutrient plays a vital role in physiological processes necessary for pathogen defense, it is often difficult to disentangle the role of an individual macronutrient on disease pathology. We explored the effects of dietary macronutrient composition on disease pathology and feeding behavior of canaries (*Serinus canaria domestica)* infected with *Mycoplasma gallisepticum* (MG). First, we provided canaries with isocaloric diets comprised of identical ingredients that varied in macronutrient content (high-protein or high-lipid) then MG- or sham-inoculated birds. We then offered both diets to canaries before and after MG- and sham-inoculation. In experiment one, high protein diet birds consumed more food than high lipid diet birds and experienced a more pronounced decrease in food intake after infection. High protein diet birds were more tolerant to MG infection, exhibiting reduced pathology when compared to high lipid diet birds, despite the two treatments having similar levels of MG-specific antibodies and MG loads. When birds had access to both diets, they consumed more of the high protein diet and experienced pathology for less time than lipid or protein restricted birds. These results highlight that macronutrient makeup of the diet can shape host tolerance and pathology and thus affect host-pathogen transmission dynamics.

**Summary Statement:** Macronutrient composition of the diet shapes tolerance in an avian host pathogen system. Birds that had greater access to protein exhibited reduced pathology despite similar pathogen loads when compared with birds with greater access to lipids.

## Introduction

Diet can be important to a host’s susceptibility to infection, severity of pathology, and ability to efficiently clear an infection (Hall et al. 2009; Unckless et al. 2015; Napier et al. 2019; Gomes et al. 2023). The effect of food availability and composition on disease dynamics within and across species has become increasingly critical to understand as more anthropogenic food resources are introduced to natural and altered environments (Becker et al. 2018). Changes in land-use and availability of resources can result in emerging disease hotspots, primarily by altering species diversity and increasing interactions within and among species (Plowright et al. 2011; Hosseini et al 2017; Becker et al. 2018). However, few studies have focused on how shifts in diet composition can influence individual disease outcomes, despite the important role individual-level disease outcomes play in epidemic dynamics (Hite and Cressler 2018).

While nutritional resources are critical to host immune function, those resources are also accessible to pathogens for growth and replication (Grieger and Kluger 1978; Cornet et al. 2014; Cressler et al. 2014). Hosts can restrict pathogen access to nutrients (i.e., starve the pathogen) by undergoing self-induced anorexia (Adamo 2006). However, hosts also require nutritional resources for mounting a robust immune response, thus a reduction in nutritional intake may have immunological or energetic costs for the host that affect disease outcomes. Researchers have proposed that relationships between food resources and infection outcomes vary across systems and contexts (Hall et al. 2009; Cressler et al. 2014; Hite and Cressler 2018; Pfenning-Butterworth 2023). In some situations, host resources are diverted and used for pathogen growth (pathogen priority hypothesis), making anorexia advantageous, and in other situations resources are used directly by the host for immune responses (immune priority hypothesis), making anorexia costly (Cressler et al. 2014). In vertebrates, abundance of nutritional resources tends to favor hosts (Cressler et al. 2014), presumably because resources for parasites or pathogens are abundant in vertebrate hosts, such that additional resources have little direct benefit to the parasites and pathogens (but see Cornet et al. 2014). Instead, additional resources simply improve host immune responses to diminish parasite and pathogen load. However, robust immune responses can also come at a cost to the host by increasing inflammation and tissue damage (Graham et al. 2005; Sorci & Faivre 2009; Long & Graham 2011). For example, one study determined that canaries infected with *Plasmodium relictum* with supplemental food had greater pathology than non-supplemented canaries, whereas parasite growth was greatest in non-supplemented canaries (Cornet et al. 2014). These studies underscore the complexity of relationships that exist between resource availability and host and pathogen outcomes.

Abundance of resources are certainly important to host-pathogen interactions, but so is the nutritional makeup of those resources. The nutritional composition of the diet contributes to how effectively immune processes eliminate pathogens (Cunningham-Rundles et al. 2005, Amar et al. 2007). Manipulative studies in an ecological context show the importance of dietary macronutrients on immune processes to ultimately affect individual disease outcomes (Klasing 2007; Cotter et al. 2011; Povey et al. 2009, 2013). In one study, caterpillars experimentally infected with a baculovirus, *Spodoptera exempta*, preferred high protein diets when given the option. The increase in consumption of high protein diets led to higher rates of survival from infection than the caterpillars given a high carbohydrate diet (Povey et al. 2013). Another study in poultry demonstrated how variation in dietary lipids affected immune responses by modifying leukocyte production (Friedman and Sklan 1995) and lipid composition can have direct effects on inflammatory responses (Klasing 1998). However, high lipid diets can increase mortality rates in invertebrates during some infections (Adamo et al. 2008). Variation in immune responses based on macronutrient composition in the diet is likely a result of a host requiring different nutrition to produce robust immune responses for clearing specific pathogens (Hite 2018). These studies demonstrate that outcomes of host-pathogen interactions depend on the macronutrient breakdown of food resources, presenting the opportunity for macronutrient specific changes in host feeding during infection and disease outcomes as seen in some invertebrate systems (Povey et al. 2013; Sieksmeyer et al. 2022). There is some evidence that vertebrates, which will selectively eat plants with antiparasitic compounds during parasitic infection (Huffman and Seifu 1989; Hutchings et al. 2003), may also shift macronutrient intake during infection as noted by reduced protein intake in zebra finches after injection with a non-pathogenic antigen (Love et al. 2024). This finding raises the possibility that vertebrate hosts could selectively consume macronutrients that improve host immune function (the immunity priority hypothesis, Cressler et al. 2014), while still reducing important macronutrients for pathogen growth (pathogen priority hypothesis; Cressler et al. 2014).

The goal of this study was to explore the effects of macronutrients on disease outcomes, e.g., disease severity and recovery time after exposure to a bacterial pathogen in a vertebrate host-pathogen system. Similar research has been done using a non-pathogenic novel antigen (Love et al. 2024); however, a true pathogenic infection could lead to different results because the immunological pathways used to clear the antigen by the host can be different (Koch et al. 2018, Macpherson et al. 2001). Often studies exploring immune and diet interactions change food abundance or food types (Tschirren et al. 2007; Hall et al. 2009; Cornet et al. 2014; Nwaogu et al. 2020), making it difficult to pinpoint the effects of macronutrients on disease outcomes. This study will improve our understanding of the role of macronutrients in shaping vertebrate disease pathology during infection and why some animals exhibit selective or reduced feeding during illness. To do this, we conducted two experiments aimed at answering different but complementary questions: 1) How will macronutrient specific diets affect disease severity and immune response in a vertebrate host-pathogen system and 2) will dietary preference shift during an infection?

To test our hypotheses, we used a common avian host-pathogen system, domestic canaries infected with the bacterial pathogen *Mycoplasma gallisepticum* (MG). MG can infect several songbird species and tends to cause severe conjunctival swelling and lethargy in several species belonging to the Family Fringilidae, including canaries and house finches (Hawley et al. 2011; Dhondt et al. 2008; Love et al. 2021). Although MG is most well-studied in the wild house finches, domestic canaries are proving to be a useful lab model species because they thrive in lab conditions and experience similar pathology and pathogen load when infected with MG as house finches (Hawley et al. 2011; Love et al. 2021; Sauer et al. 2024). Earlier studies in our lab indicated that zebra finches reduce protein consumption but maintain lipid consumption during a non-pathogenic immune challenge (Love et al. 2024), suggesting that lipids may benefit host immune responses. Hosts may reduce protein consumption because protein is more likely to contain iron than other macronutrients and iron is essential for some pathogens (Kluger & Rothenburg 1979; Cassat & Skaar 2013). Thus, reducing protein consumption during an immune threat could limit pathogen growth. Based on these data and the immunity and pathogen priority hypotheses, we predicted in the first experiment, that birds fed a high lipid diet would clear MG faster and have lower pathogen load than birds fed the high protein diet. Because of these benefits to the host, we predicted that birds in the second experiment would prefer a high lipid diet during MG infection.

## MATERIALS AND METHODS

### Experimental Design and Timeline

#### Experiment 1

Individually housed female canaries (N=37) were provided either a high lipid (n=22) or high protein (n=20) diet 17 days prior to inoculation to acclimate treatment groups to the diets. Following acclimation to the diets, birds were inoculated with either FREY’s media or MG, resulting in 4 treatments: Lipid MG (n = 13), Lipid control (n = 9), Protein MG (n = 10), Protein control (n = 10). Two birds in the Lipid MG treatment and one in the Protein MG treatment died during the experiment, so they are not represented at all time points in the dataset. Only females were used in this experiment because they were being used in a subsequent study exploring disease-mediated parental effects on egg attributes. We weighed birds, measured fat scores, assessed eye inflammation, and measured various immune endpoints (e.g. white blood cell counts, pathogen load, MG-specific antibody levels) throughout infection. Eyes were scored for inflammation and swabbed to measure pathogen load prior to infection and every other day post inoculation until 35 days post infection. Body mass and fat scores were collected prior to infection and at days 7-, 14-, 21- and 35-days post infection. We collected blood samples from birds immediately prior to infection and at days 7-, 14-, and 21-days post infection. The blood samples were used to assess hematocrit and white blood cell differential counts, and to quantify MG antibody concentrations in birds.

#### Experiment 2

Canaries (N=25) were housed individually and provided both a high lipid and high protein diet daily throughout the experiment. Birds were acclimated to the diet for 17 days, then inoculated with either FREY’s media (females: n = 6, males: n = 6) or MG (females n = 6; males n = 7). A mix of males and females were used in this experiment because we did not have enough females to keep this consistent with the design of experiment 1. Four MG-infected birds died during the first week after MG-inoculation, three males and one female. We recorded bird’s body mass, fat stores, conjunctiva inflammation, and other immune endpoints (e.g. white blood cell counts, pathogen load, MG-specific antibody levels) throughout infection. Prior to and every 2-3 days after infection, we scored eye inflammation, then bilaterally swabbed eye conjunctiva to quantify pathogen load. Body mass and fat scores were collected prior to infection and 7-, 14-, 21-, and 35 days post infection. We collected blood samples prior to infection and at days 7-, 14-, and 21-days post infection. Like experiment 1, blood samples were used to assess hematocrit, determine relative abundance of white blood cells, and quantify concentrations of MG-specific antibodies.

### Bird Housing

Canaries were housed in an ABSL-1 biosafety room on a 14L:10D light cycle throughout both experiments. Birds were housed in wire cages (24”x16”x16”) that are divided into two units with each unit containing one bird to allow assessment of individual food consumption. Each housing space contained two plastic perches, a water dish, and a food dish in the first experiment. The housing space contained an additional food dish for the second experiment (diet preference) and the water dish was placed completely within the cage. To prevent contamination of control birds by MG-infected birds, which is primarily transmitted through direct contact and fomites in finches (Dhondt et al. 2007), a plastic partition divided the room with controls held on one side of the partition and MG-infected birds kept on the other.

### Diet composition and monitoring feeding

The diets we fed to birds were chosen to allow birds access to all macronutrients and create a scenario that would allow birds to distinguish between the two diets when they were offered together (experiment 2) but would prevent canaries from making up for the reduced proteins or lipids by consuming more of the diet they were given (experiment 1). In the first experiment, birds were fed daily either a 24 g isocaloric food bar that was lipid-rich (80:20 lipid to protein ratio) or protein-rich (20:80 lipid to protein ratio). Both diets contained varying proportions of egg whites, egg yolks, hulled millet, cod liver oil, and were congealed together with agar. In the second experiment, birds received a bar of each diet placed in separate food dishes. The dishes were on either side of the cage front, and we randomized which side received which diet for each rack of cages at the beginning of the experiment. Diet placement did not change during the experiment. Diets were weighed and replaced daily shortly after lights came on. For both experiments, three food bars for both diets were placed in the bird room and weighed daily to account for desiccation, which were then averaged and subtracted from the amount of diet consumed by each bird.

### Inoculations and monitoring of disease severity

After acclimation to diets, birds were inoculated with MG or a control solution. We inoculated MG-treated birds in both experiments bilaterally with MG inoculum (VA1994; E. Tulman, University of Connecticut) in their palpebral conjunctiva with 25 μL containing 5*107 CCU/ml of MG inoculum diluted 16.9% in Frey’s media. Birds assigned to the control treatments in both experiments were inoculated with 25 μL of Frey’s media. Throughout the duration of infection, inflammation of the conjunctiva, a measure of disease severity, was scored on a scale of 0-3 (Hawley et al. 2011); higher scores represent a greater degree of disease pathology. Both eyes of each bird were given a score and were summed to determine a total eye score value (Hawley et al. 2011). Disease severity was also monitored by recording changes in body mass and fat scores. Fat scores were measured on a scale of 0-3 and categorized by how much visible adipose tissue was present in the interclavicular fossa of the birds. A low fat score value indicates only a small trace or lack of visible fat tissue and higher values indicate more fat tissue present.

### Antibody Assays

After blood samples were collected, they were microcentrifuged at 3500 rpm, the plasma was removed and stored in a −20°C freezer for future analysis of MG antibody concentrations. Serum antibodies were quantified using the IDEXX MG antibody enzyme-linked immunosorbent assay test kit (IDEXX, Cat#99-06729). A blocking step was added to the original assay kit’s protocol, with the addition of 300 μL of 1% bovine serum albumin (Pierce 10X BSA; Thermo Fisher Scientific) in phosphate-buffered saline to room temperature plates before they were incubated. All plates were washed three times with phosphate-buffered saline containing 0.05% Tween 20 using an ELx50 plate washer (BioTek). Serum samples were diluted 1:50 in sample buffer and were then plated to be run in duplicate. Intensity of light absorbed by the serum samples was measured at 630 nm using a spectrophotometer and an ELISA value was then calculated.

### Pathogen load

We determined pathogen load using quantitative PCR following the procedure outlined by (Grodio et al. 2008) that targets the mgc2 gene of MG. Sterile cotton swabs were dipped in tryptose phosphate broth and used to swab conjunctiva in both eyes of birds for five seconds each. The tips of the swabs were then cut off and placed in 300 μl of the tryptose phosphate broth and frozen in a −20°°C freezer. Qiagen DNeasy 96 Blood and Tissue kits (Qiagen, Valencia, CA) and primers and probe that target mgc2 were used to extract genomic DNA. The total liquid volume of 15 μl included 7.5 μl of Primetime Master Mix (Bio-Rad Laboratories, Hercules, CA), 3.525 μl DNase-free water, 3 μl of DNA sample, 0.375 μl of forward and reverse primers and 0.225 μl of 10 um MG probe. A BioRad CFX-96 machine was used for cycling at 95°C for three minutes, followed by 40 cycles of 95°C for three seconds, and then 60°C for 3 seconds. The ramp rate of the machine was set to 0.5/second. The mgc2 values were a summed total of both conjunctiva of each individual bird and sample day. Final concentrations were calculated by multiplying 3 μl, the amount of DNA sample used, by 66.666, to be comparable to the 200 μl that was produced from the elution step. Determined values for mgc2 were then log transformed to reduce the large outputs that were generated.

### Statistical Analyses

Prior to all statistical analyses, data were checked for normality and homoscedasticity. All statistics were conducted with R version 4.3.2 in R Studio (RStudio Team, 2021). To test for the effects of diet, MG exposure, time, and their interactions in the first experiment and sex, MG exposure, time, and their interactions in the second experiment on food intake (g), body mass (g), fat scores, and hematocrit (%), we conducted linear or normally distributed generalized linear mixed-effects models followed by ANOVAs (*lme4*, *glmmTMB*, & *car* packages). To test for effects of the same predictors on relative white blood cell abundance and heterophil:lymphocyte ratio, we conducted separate generalized linear mixed-effects models with varying distributions followed by ANOVAs (see Tables S9 & S19 for details). To test for the effect of diet, time, and their interaction in the first experiment and sex, time, and their interaction in the second experiment on total eye score in MG-exposed birds, we conducted a Poisson distributed generalized additive mixed model that included a smoothing spline to model the non-linear effects over time (*mgcv* package). We further explored the effects of diet from the first experiment on pathology recovery time (days) by conducting a normally distributed generalized linear model followed by an ANOVA. To test for the effects of diet, time, and their interaction in the first experiment and sex, time, and their interaction in the second experiment on log10-transformed pathogen load and MG-specific antibody level (optical density), we conduced normally distributed generalized linear mixed-effects models followed by ANOVAs. All mixed-effects models included a random intercept for bird identity. In both experiments, we analyzed food consumption two ways, by day and by week, which yielded similar results. We chose to use weekly feeding patterns because they were easier to compare visually, and capture feeding behavior at distinct phases of infection: pre-infection, peak infection, early recovery, and late recovery. See supplemental materials for daily feeding patterns (Figures S1 & S3).

## Results

### Experiment one

High protein birds consumed more food each week than high lipid birds, except in the week immediately following inoculation when food consumption decreased sharply for the high protein birds and moderately for the high lipid birds (infection*diet*week: *X ^2^* = 5.10, *df* = 1, *p* = 0.02; diet*week: *X ^2^* = 5.97, *df* = 1, *p* = 0.01; diet: β_protein_ = 7.31±9.98, *X ^2^* = 3.30, *df* = 1, *p* = 0.07; week: β = 0.74±1.10, *X ^2^* = 2.80, *df* = 1, *p* = 0.09; Figure 1A & Table S1). There were no other significant main or interactive effects on food consumed (*p* > 0.1; Table S1).

**Figure 1.**
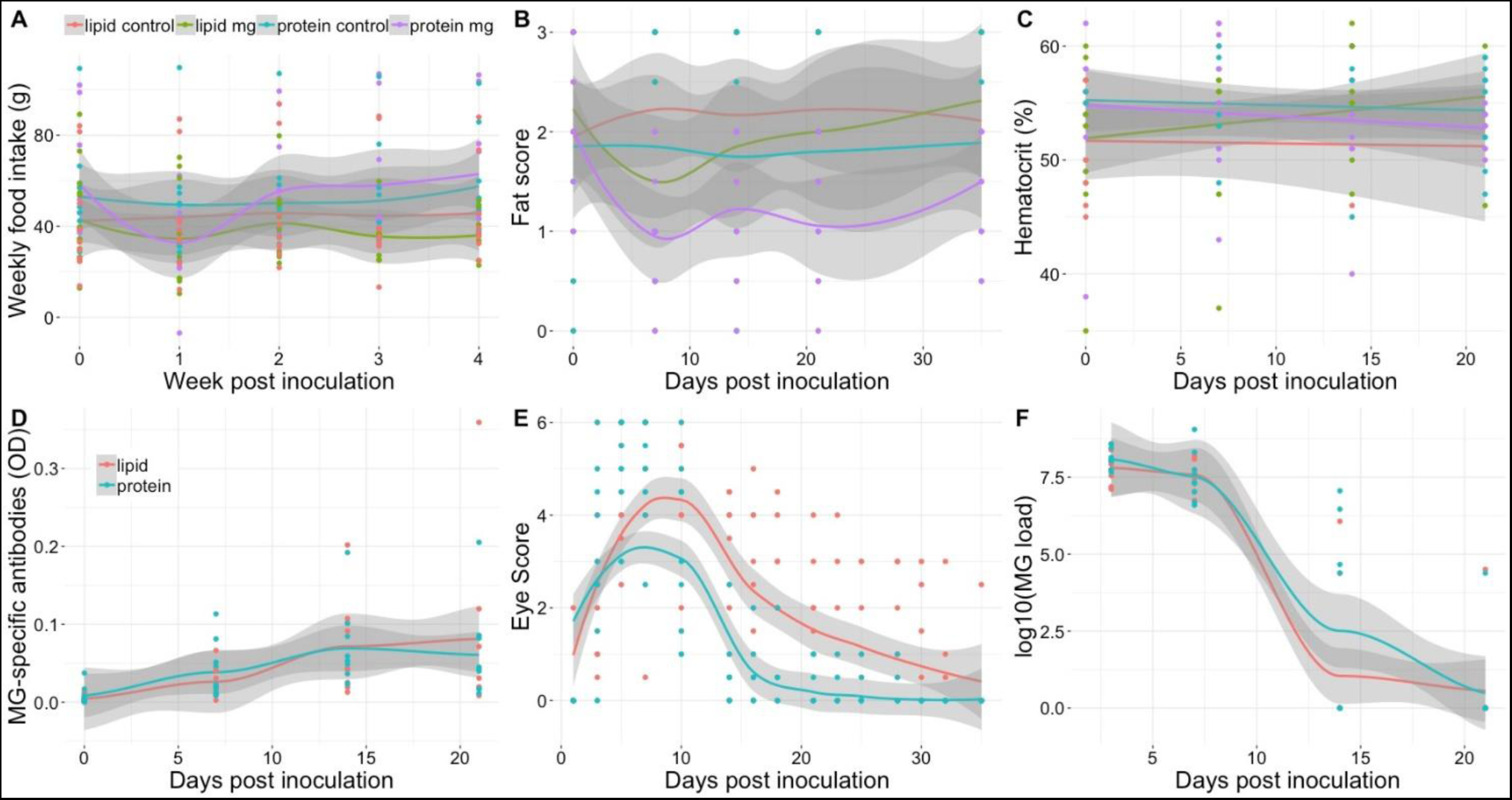
Weekly food intake (g) (A), furcular fat scores (B), % hematocrit levels (C), MG-specific antibody production (D), conjunctival swelling represented by a score of swelling severity (E), and MG pathogen load (F) measured in female canaries (Serinus canaria domestica) that were fed an isocaloric diet that differed in macronutrient ratios and were later inoculated with Mycoplasma gallisepticum (MG) or Frey’s media (controls). Birds were inoculated seventeen days after receiving either the high protein (20:80 lipid:protein) or high lipid (80:20 lipid:protein) which was termed day 0. Birds remained on the diet for the duration of the experiment. Only MG-infected birds are depicted in figures related to disease outcomes (antibody production, eye swelling, and pathogen load). Lipid control (n = 13); lipid MG (n = 9); protein control (n = 10); protein mg (n = 10). Points represent raw data with average trend lines surrounded by 95% confidence interval bands in grey.

Regardless of treatment, all birds were similar in mass (Figure S1) and had similar furcular fat scores (Figure 1B) prior to MG infection. Body mass decreased during the middle of the experiment, then rebounded, but was not affected by infection or diet (day: β = −0.03±0.02, *X ^2^* = 3.13, *df* = 1, *p* = 0.08; Table S2). Hematocrit differed over time by diet with higher hematocrit in birds fed the protein diet at the start of the experiment, but lipid birds catch-up by the end of the experiment (day*diet: *X ^2^* = 5.31, *df* = 1, *p* = 0.02; Table S3). There were no other significant main or interactive effects on body mass, fat scores or hematocrit (*p* > 0.1; Tables S2-S4).

There was a significant effect of time on eye swelling (day: *F* = 130.6, *edf* = 1.99, *p* < 0.0001; Figure 1E & Table S5), in which MG-infected birds from both diets exhibited more severe swelling during the first 10 days of infection and swelling diminished in the days after peak infection (Day: 7-10). Macronutrient composition of the diet also significantly affected eye score, with more severe swelling in the birds fed the high lipid diet as compared to birds fed the high protein diet (Diet: *t* = −2.90, *p* = 0.004; Figure 1E & Table S5). On average the time to recovery of conjunctiva swelling of birds was 16.9 ± 2.4 days in MG infected birds fed the high protein diet and 25.7 ± 3.4 days for MG infected birds fed the high lipid diet (β_protein_ = −8.78±4.29, *X ^2^* = 4.19, *df* = 1, *p* = 0.04; Table S6).

Infected birds in both diet treatments exhibited high pathogen loads in the first week of infection, which decreased by 14 days post infection (β = −0.41±0.04, *X ^2^* = 171.05, *df* = 1, *p* < 0.001; Figure 1F & Table S7) and did not differ with diet treatment (*p* > 0.4; Table S7). Similarly, infected birds in both diet treatments experienced a significant increase in MG specific antibodies after infection (β = 0.01±0.001, *X ^2^*= 25.93, *df* = 1, *p* < 0.001; Figure 1D & Table S8) and did not differ among diets (*p* > 0.3; Table S8).

The series of repeated measures ANOVAs revealed significant effects of infection and diet on production of different white blood cell types. In general, basophils remained low throughout the entire experiment, with individuals exhibiting 0-1 basophils at most time points. However, because one individual in the MG protein treatment had five basophils on day 14 post inoculation (all other birds in this treatment on day 14 had 0-2 basophils) this generated a significant infection*diet*day interaction on basophil production (*X ^2^*= 4.65, *df* = 1, *p* = 0.03; Figure S2 & Table S9). In all groups, eosinophils increased from the day of inoculations to day 7 post inoculation and remained elevated to day 21 post inoculation (β = 0.04±0.02, *X ^2^* = 24.64, *df* = 1, *p* < 0.001; Figure S2 & Table S9) and lipid birds had more eosinophils than protein birds (β_protein_ = −0.51±0.42, *X ^2^*= 4.04, *df* = 1, *p* < 0.05; Figure S2 & Table S9). Monocytes trended towards increasing throughout the course of the experiment in all birds (β = 0.04±0.03, *X ^2^* = 3.14, *df* = 1, *p* = 0.08; Figure S2 & Table S9). On average birds fed the high protein diet experienced significantly increased numbers of lymphocytes (β_protein_ = 4.07±3.19, *X ^2^*= 9.80, *df* = 1, *p* = 0.002; Figure S2 & Table S9) as compared to birds fed the high lipid diet. Lymphocytes also generally decreased in abundance over time (β = −0.59±0.18, *X ^2^* = 24.80, *df* = 1, *p* < 0.001; Figure S2 & Table S9). Heterophil production increased slightly with time (β = 0.01±0.01, *X ^2^* = 8.61, *df* = 1, *p* = 0.003; Figure S2 & Table S9) and protein birds trended towards lower abundance of heterophils than lipid birds (β_protein_ = −0.51±0.26, *X ^2^* = 3.55, *df* = 1, *p* = 0.06; Figure S2 & Table S9). The heterophil:lymphocyte ratio was not affected by infection status of birds but did increase with time across all treatments (β = 0.03±0.01, *X ^2^* = 40.62, *df* = 1, *p* < 0.001; Figure S2 & Table S9) and tended to be lower in protein birds than lipid birds (β_protein_ = −0.55±0.26, *X ^2^* = 4.95, *df* = 1, *p* = 0.03; Figure S2 & Table S9). There were no other significant main or interactive effects on white blood cell relative abundance or heterophil:lymphocyte ratios (*p* > 0.1; Table S9).

### Experiment two

Infected birds in experiment two ate less food than control birds and all birds increased food intake over time (week: β = 10.17±1.87, *X ^2^*= 77.37, *df* = 1, *p* < 0.001; infection: β_mg_ = −17.12±10.41, *X ^2^*= 6.66, *df* = 1, *p* = 0.01; Figure 2A & Table S10). We see the same pattern in feeding when we compare intake of the different diets; infected birds ate less protein diet (β_mg_ = −0.03±0.25, *X ^2^* = 5.77, *df* = 1, *p* = 0.02; Figure 2B & Table S11) and lipid diet during the experiment than control birds (β_mg_ = −3.28±6.15, *X ^2^* = 3.75, *df* = 1, *p* = 0.05; Figure 2C & Table S12), and all birds increased protein and lipid intake over time (protein: β = −0.03±0.01, *X ^2^* = 26.82, *df* = 1, *p* < 0.001; lipid: β = 6.48±1.26, *X ^2^* = 57.50, *df* = 1, *p* < 0.001). We also found that females tended to consume more food overall and more protein each week than males (total: *X ^2^* = 2.64, *df* = 1, *p* = 0.10; protein: *X ^2^* = 5.77, *df* = 1, *p* = 0.02). There were no other main or interactive effects of sex on canary feeding patterns (*X ^2^*< 2.5 & *p* > 0.11).

**Figure 2.**
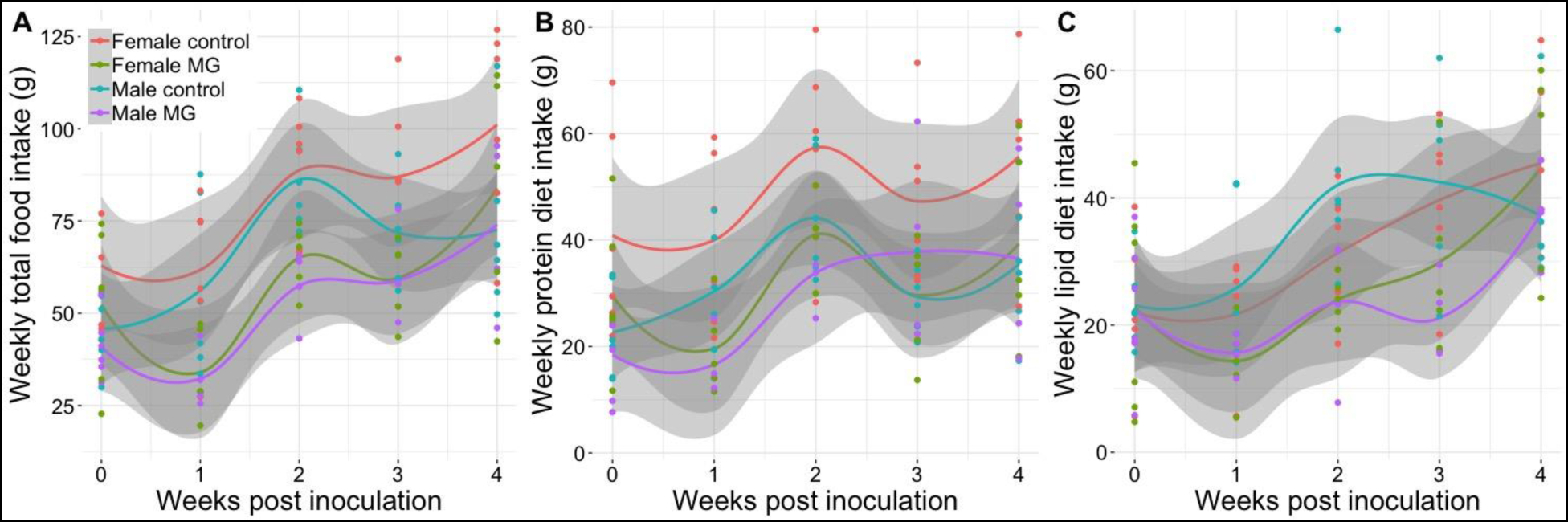
Weekly food intake (g) of male and female canaries (*Serinus canaria domestica*) provided with two isocaloric diet bars, one that was high in protein (20:80 lipid:protein) and one that was high in lipid (80:20 lipid:protein) for seventeen days before inoculation with *Mycoplasma gallisepticum* (MG) or Frey’s media (controls). The day of inoculation was termed day 0. Weekly total food intake (A), consumption of the high protein bar (B), and consumption of the high lipid diet bar (C) was monitored for four weeks after inoculation. Controls: females: n = 6, males: n = 6; MG-infected canaries: females n = 6; males n = 7. Points represent raw data with average trend lines surrounded by 95% confidence interval bands in grey.

Birds infected with MG weighed less than control birds (β_mg_ = −4.57±2.35, *X ^2^* = 4.04, *df* = 1, *p* = 0.04; Figure 3B & Table S13) and had lower hematocrit (β_mg_ = −0.85±3.08, *X ^2^* = 4.25, *df* = 1, *p* = 0.04; Figure 3C & Table S14). Although females were heavier than males (β_male_ = −5.09±2.34, *X ^2^* = 3.81, *df* = 1, *p* = 0.05), males had higher hematocrit than females (β_male_ = 8.78±3.05, *X ^2^* = 10.26, *df* = 1, *p* = 0.001). Infected birds lost fat stores later in infection, whereas control birds tended to gain fat stores (infection*day: *X ^2^* = 3.81, *df* = 1, *p* = 0.05; Figure 3A & Table S15). Females also had more fat stores than males at the beginning of the experiment, but males gained fat later in the experiment (sex: β_male_ = −1.10±0.40, *X ^2^* = 6.43, *df* = 1, *p* = 0.01; sex*day: *X ^2^*= 7.48, *df* = 1, *p* = 0.006). There were no other significant main or interactive effects on body mass, fat stores, or hematocrit of canaries (*p* > 0.10).

**Figure 3.**
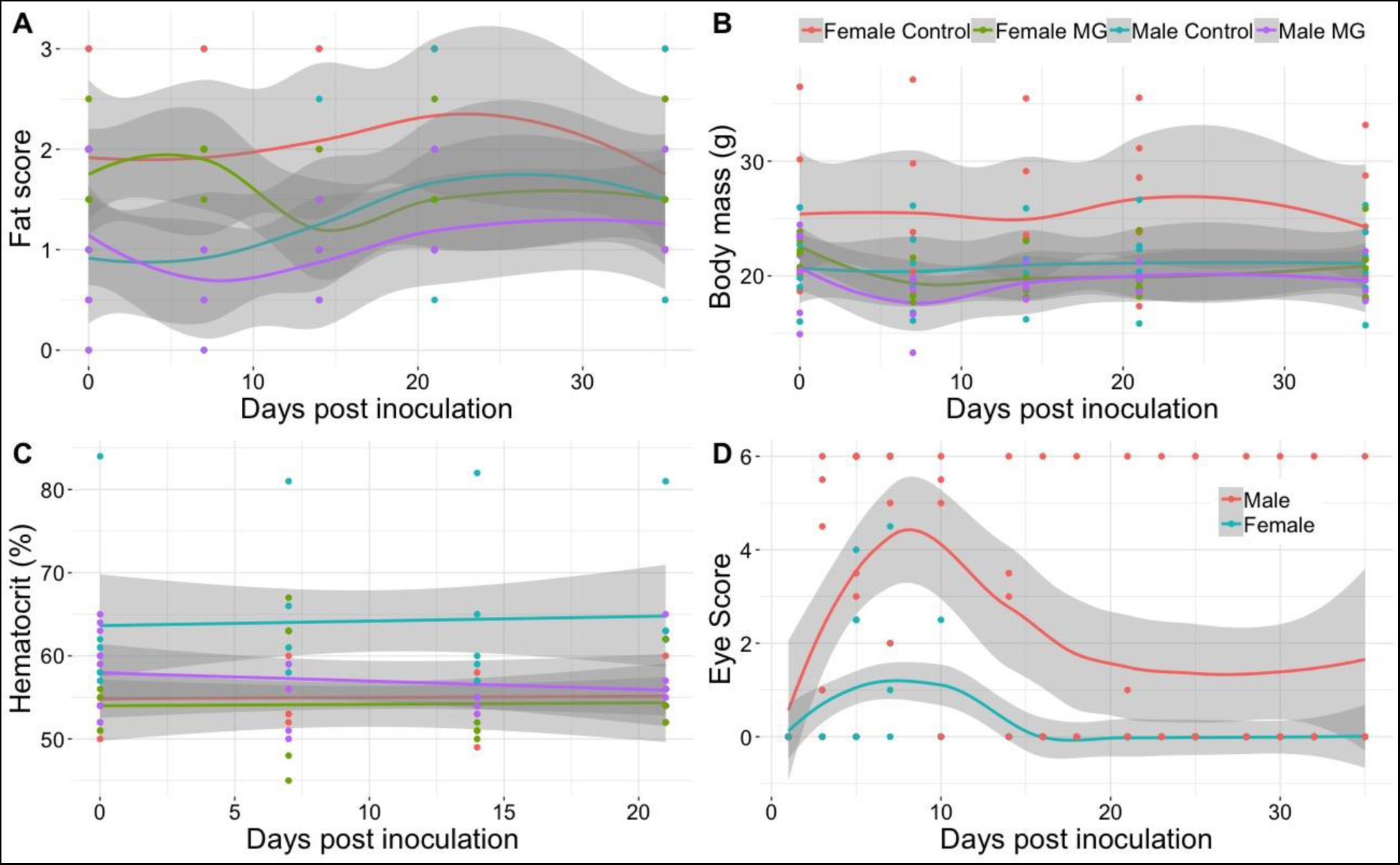
Furcular fat stores (A), body mass (B), % hematocrit (C) and conjuctival swelling represented by a score of swelling severity in male and female canaries (*Serinus canaria domestica*) provided with two isocaloric diet bars, one that was high in protein (20:80 lipid:protein) and one that was high in lipid (80:20 lipid:protein) for seventeen days before inoculation with *Mycoplasma gallisepticum* (MG) or Frey’s media (controls). The day of inoculation was termed day 0 and endpoints were monitored for four weeks after inoculation. Only MG-infected birds are depicted in the figure related to disease outcomes (eye swelling). Controls: females: n = 6, males: n = 6; MG-infected canaries: females n = 6; males n = 7. Points represent raw data with average trend lines surrounded by 95% confidence interval bands in grey.

Eye swelling, pathogen load, and MG antibodies increased in infected birds after inoculation with MG. Swelling and pathogen load was greatest 5-10 days after infection, then subsided (*F* = 3.964, *edf* = 1.84, *p* = 0.07; Figure 3C & Table S16), whereas MG antibodies steadily increased after inoculation (β = 0.001±0.001, *X ^2^* = 18.02, *df* = 1, *p* < 0.001; Figure S4 & Table S18). Infected males exhibited greater swelling (*t* = 8.73, *p* < 0.001) and experienced greater pathogen loads (β_male_ = 1.35±1.89, *X ^2^* = 4.59, *df* = 1, *p* = 0.03; Figure S4 & Table S17) during infection. Infected males also produced more MG-specific antibodies over time than females (sex*day: *X ^2^*= 3.56, *df* = 1, *p* = 0.06). Aside from one bird that exhibited swollen conjunctiva past the end of the second experiment, infected birds recovered by day 16 post infection and average recovery time was 11.2 ± 3.0 days.

The only effect of MG infection on the relative abundance of white blood cells in birds that had both diets available was dependent on bird sex. Monocytes decreased in male birds over time with a peak occurring at day 7 post infection only in MG-infected males, which aligns with peak pathology, while monocytes in female birds stayed relatively consistent over time with MG-infected female birds having lower levels than control females (sex*day: *X ^2^* = 4.52, *df* = 1, *p* = 0.03; sex*infection: *X ^2^* = 4.37, *df* = 1, *p* = 0.04; Figure S4 & Table S17). Heterophils increased slightly for female birds over time, regardless of MG exposure, but stayed consistent over time in male birds (sex*day: *X ^2^* = 4.41, *df* = 1, *p* = 0.04; Figure S4 & Table S17). Heterophil:lymphocyte ratios varied as a function of sex and time where it slightly increased for females over time but slightly decreased for males (sex*day: *X ^2^* = 13.61, *df* = 1, *p* < 0.001; Figure S4 & Table S17). Similar to experiment one, the presence of basophils was rare and only one was detected throughout the course of this experiment. There was no significant difference detected in relative abundance of any other white blood cell between control and MG-infected birds or between sexes (*p* > 0.09).

## Discussion

In this study, we used a vertebrate host-pathogen system to shed light on how individual macronutrients affect host disease outcomes and if these effects are reflected in dietary preference during an infection. Our results indicate that a diet rich in lipids but poor in proteins results in greater disease pathology and reduced recovery time despite similar infection intensity and antibody production (i.e., reduced tolerance) than birds receiving a diet rich in protein but poor in lipids. However, when given the choice, infected birds do not selectively reduce lipids following infection, although, the birds do consume more protein overall regardless of infection status. Our results support the immune priority hypothesis, because macronutrients shaped host immune responses but not pathogen growth. Our results also demonstrate that diet quality has implications for disease outcomes important for transmission and reveals an immunological mechanism through which altered food landscapes, whether human-induced (e.g., dumpsters, bird feeders, etc.) or naturally occurring (e.g., drought), can alter wildlife disease dynamics.

A high protein diet was important to immune support noted by the faster attenuation of disease pathology as measured by conjunctival swelling (Hawley et al. 2011) in high protein birds than high lipid birds, though both groups exhibited similar peak pathology. Conjunctival swelling is an important indicator of transmission in avian MG systems. In house finches, conjunctival swelling correlates positively with conjunctival pathogen load and the likelihood of transmitting MG (Adelman et al. 2013). However, in our system, pathogen load and MG antibody production did not differ between birds fed the two diets, despite the extended pathology in the high lipid birds. This finding has several important implications. First, high lipid birds are more likely to transmit MG because swelling alone is integral to pathogen transmission, perhaps because they are more likely to wipe infected tissues onto feeders or conspecifics (Adelman et al. 2013). However, high protein diet birds also may be capable of transmitting MG even though the conjunctiva is not heavily swollen due to high pathogen load. This could alter typical transmission dynamics in a population because birds can detect illness in conspecifics (Bouwman and Hawley 2010; Love et al. 2021; Love et al. 2023) and engage in avoidance behaviors to prevent illness, but if an important indicator of illness (e.g., swelling) is missing they may not detect illness in birds eating a protein rich diet even though they are capable of transmitting the pathogen. Second, high protein birds appear to tolerate infection better than high lipid birds because they reduced pathology despite having similar pathogen loads and antibody production to birds fed a high lipid diet. This indicates that diet plays an important role in whether individuals tolerate or resist an infection and suggests that studies exploring how resource availability shapes host-pathogen evolutionary dynamics are needed.

The differences in tolerance and pathology driven by diet that we detected are also relevant to consider in the context of supplemental feeding of wildlife as studies have shown that when supplemented foods are not carefully considered they can lead to global scale shifts in pathogen transmission (Murray et al. 2016). Resource driven shifts in immunity can drastically alter epidemics by increasing the likelihood of producing individuals with superspreading abilities, because resource scarcity is more likely to result in hosts with high pathogen loads or extended infections (Hall 2019). The longer period of conjunctival swelling in the high lipid birds in our study, which experienced protein scarcity, should be more likely to transmit MG than the high protein birds (Sauer et al. 2024).

Although the mechanism behind the faster recovery in high protein birds is unclear, this result could suggest that a certain threshold of protein intake is required prior to and during infection for optimal recovery. Dietary protein is essential for the regulation and activation of both T and B lymphocytes, both of which are important for recovery from an infection (Li et al. 2007) and some animals are more likely to survive infection when on a high protein diet (Jahanian 2009, Lee et al. 2008). We found that high protein birds had greater circulating lymphocytes than high lipid birds. B lymphocytes produce antibodies (LeBien and Tedder 2008); however, we did not detect a difference in MG antibody production. It is possible that dietary protein may be affecting other immunological pathways that were not investigated, like cytokines which are produced by T lymphocytes. For instance, increased levels of interleukin-1 act as pro-inflammatory agents (Dinarello 1997), while other cytokines, such as interleukin-5 and interleukin-10, play an anti-inflammatory role during inflammatory infections (Pirola and Ferraz 2017). Our microscopy techniques did not distinguish between types of lymphocytes, so it is possible that elevated T lymphocytes, but not B lymphcytes, drove the diet effect on relative abundance of circulating lymphocytes and led to the production of anti-inflammatory cytokines. We also found that Eosinophils were higher in high lipid birds. Eosinophils move into inflamed areas and cause inflammation and tissue damage (Behm and Ovington 2000) and could be contributing to increased conjunctival swelling in high lipid birds. High protein birds also had higher hematocrit at the beginning of the experiment, which can indicate greater oxygen carrying capacity, but it is unclear how this might affect pathology. Although we found differences in the underlying immune cells important in responding to an infection, more work is needed to determine the underlying immune mechanisms stimulated by diet that lead to differences in disease outcomes.

There was some evidence that it was disadvantageous to only have access to a high protein low lipid diet. Despite high protein birds clearing conjunctival swelling more quickly than high lipid birds, high protein birds had somewhat lower furcular fat stores at the end of the study (Fig 1C) as compared to high lipid birds. Scoring fat stores is a coarse measure of available fat to the bird, which may have clouded our ability to detect a significant effect of diet on this endpoint. If high protein birds did suffer greater total body fat loss this suggests that high protein birds, which consumed fewer grams of lipids than high lipid birds (0.31 ± 0.01g vs 0.69 ± 0.02g), relied heavily on fat stores to meet the energy demands of fighting the infection and may be an important cost for clearing the pathogen. We did detect significant fat loss in infected birds with access to both diets (experiment 2), but these birds quickly recovered fat stores by the end of the experiment. Infected birds with access to both diets also recovered faster from conjunctiva swelling (11d) than both the high protein (16d) and high lipid birds (25d) in experiment one. These results suggest that having access to both diets allowed infected birds to mitigate the more severe disease outcomes that resulted from consuming the macronutrient deficient diets and there are optimal thresholds for both macronutrients that must be met to both maintain body condition and tolerate infection.

In both experiments, we found that infected birds had lower food intake as compared with controls, a behavioral pattern referred to as illness-induced anorexia, which is commonly observed across vertebrates in response to pathogenic infection (Murray and Murray 1979, Adelman & Martin 2009). When birds had access to only one diet, reduced feeding was greater in high protein birds than high lipid birds (39.11% vs 17.34%), a result consistent with other research in our lab on zebra finches which reduced protein consumption following an immune challenge with the bacterial endotoxin lipopolysaccharide (Love et al. 2024). In our study, infected birds may have reduced protein intake during infection because of its importance to the pathogen (Kluger & Rothenburg 1979; Cassat & Skaar 2013), particularly when they have had ample access to dietary proteins prior to infection (Lin et al. 2020). While we expected macronutrient selective feeding following infection when birds had access to both diets (experiment 2), this did not occur. This result is counter to findings in zebra finches and caterpillars, which both exhibited macronutrient specific shifts in protein feeding after an immune challenge (reduced protein intake; Love et al. 2024) or infection (increased protein intake; Povey et al. 2009). What we may have detected in canaries is a threshold requirement of proteins. Generally, birds preferred to eat the high protein diet and the high lipid birds in experiment one already had lower protein intake (0.67 ± 0.01g vs 2.97 ± 0.07g), therefore they did not need to or could not reduce protein intake much further. Regardless, these data indicate that birds can discern differences in macronutrient make-up of the diet and feed in ways that maximize immunity, but they may not be able to shift feeding in response to an infection aside from general illness-induced anorexia. Future studies should explore whether birds can exhibit prophylactic shifts in feeding behaviors in response to heightened infection risk (Love et al. 2024).

In the second experiment, we detected several sex effects on the behavioral and immunological endpoints that we measured. Our study was not focused on sex differences in immune responses, and males were included for logistical reasons, but the findings warrant some discussion. Females ate more food and more protein, but not lipids than males during the experiment. Perhaps, as a result, females weighed more and had more fat stores at the beginning of the experiment than males. Females also experienced lower pathology, pathogen growth, and MG antibody production, but males had higher hematocrit. It is interesting that the sex that consumed more protein also had the lowest disease pathology and greatest tolerance, which is consistent with the patterns we detected in the high protein birds in experiment one. It is still unclear what mechanism may drive this. Differences in circulating levels of eosinophils and lymphocytes did not exist between the sexes like they did between diet treatments, though males did have a larger monocyte response 7 days after infection and monocytes contribute to local and systemic inflammation (Kurihara et al. 1997). Although the sex-driven changes in white blood cells are different from diet-driven changes, in both instances the groups expressing the greatest pathology had greater relative abundance of white blood cells associated with inflammation. Importantly, the sex-specific differences in pathology and tolerance we detected can lead to sex-biased disease dynamics (Sauer et al. 2023).

This study broadens our understanding of the dynamics between diet and a host’s response to infection with a replicating pathogen. Our findings indicate that consuming higher levels of dietary protein is important for clearing the physical symptoms of an infectious pathogen by increasing host tolerance but not host resistance. These findings are important because it demonstrates that the food landscape can shape individual health outcomes in response to infection and how long individuals can transmit the pathogen. These results should be considered when predicting how anthropogenic food sources and shifts in natural food sources caused by climate change will affect wildlife epidemic dynamics.

## Acknowledgements

We thank M. Sudnick and W. Kirkpatrick for their support in the lab.

## Funding

Funds were provided by grants to S.E.D. from the National Science Foundation (1941861) and Arkansas Biosciences Institute.

## Author Contributions

SED conceived of the experiment. WGP, SED, ALC, and ELS designed the experiment, analyzed the data and wrote the manuscript. All participants collected the data.

## Ethics

All procedures were approved by the University of Arkansas Institute of Animal Care and Use Committee.

## Data availability statement

The data supporting the results are archived on GitHub: https://github.com/erinsauer/Perrine-et-al-MG-diet

## Competing interests

The authors have no competing interests to declare.

## Declaration of AI Use

AI technology was not used in the development of this manuscript or project.

## Notes

https://github.com/erinsauer/Perrine-et-al-MG-diet

### Competing Interest Statement

The authors have declared no competing interest.

