## Supplemental files for "A high lipid diet leads to greater pathology and lower tolerance during infection"

### Supplementary results for: A high lipid diet leads to greater pathology and lower tolerance during infection

Figure S1 | Effect of diet (lipid or protein) and *Mycoplasma gallisepticum* (MG) or sham inoculation on A) daily food intake (g) and B) body mass (g) of canaries (*Serinus canaria domestica*) over time in experiment one. Points represent raw data with average trend lines surrounded by 95% confidence interval bands in grey.

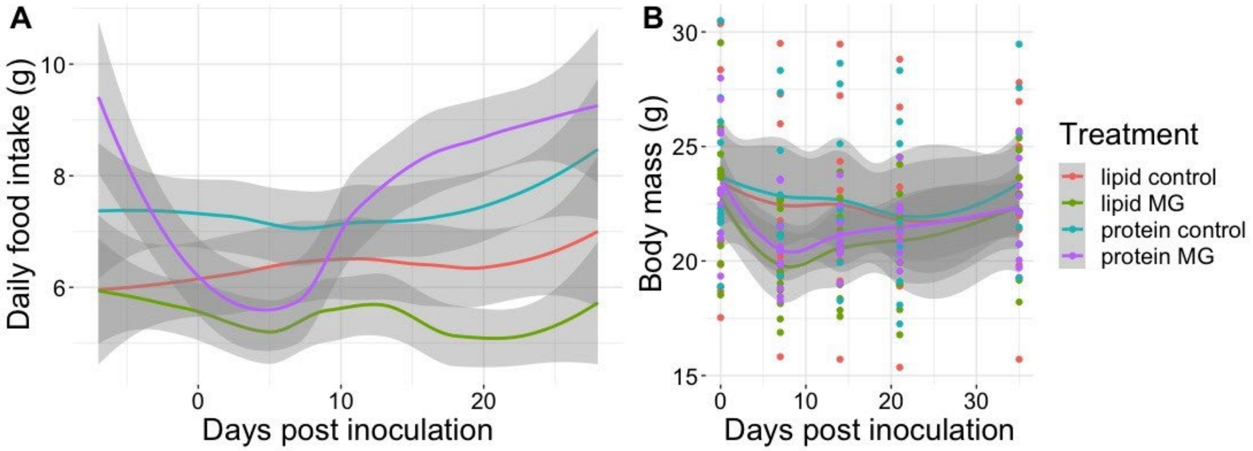

Figure S2 | Effect of diet (lipid or protein) and *Mycoplasma gallisepticum* (MG) or sham inoculation on relative abundance of white blood cells over time from canaries (*Serinus canaria domestica*) in experiment one. Points represent raw data with average trend lines surrounded by 95% confidence interval bands in grey.

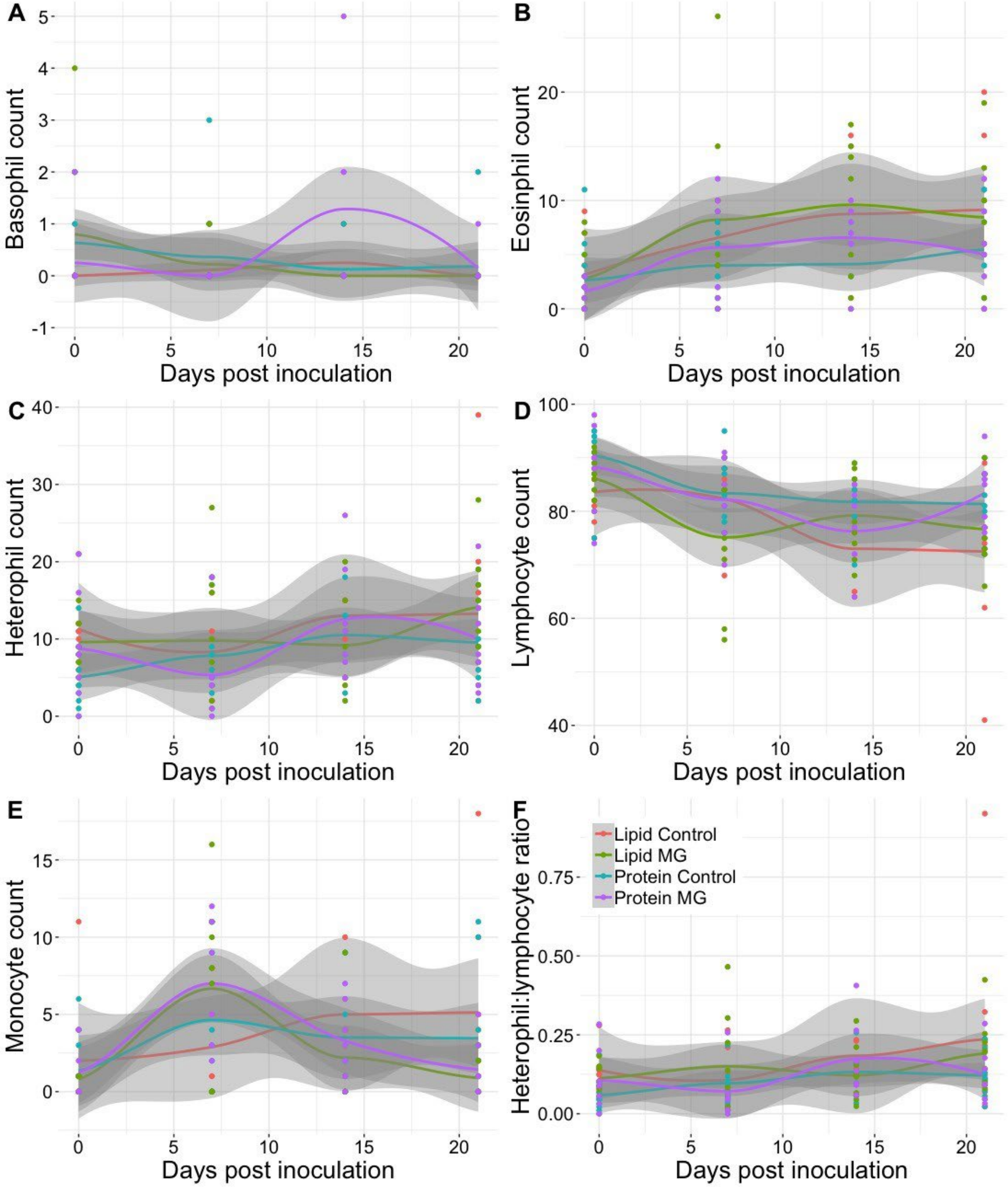

Figure S3 | Effect of sex and *Mycoplasma gallisepticum* (MG) or sham inoculation on A) total, B) protein, and C) lipid daily food of canaries (*Serinus canaria domestica*) intake over time during experiment two. Average trend lines are surrounded by 95% confidence interval bands in grey.

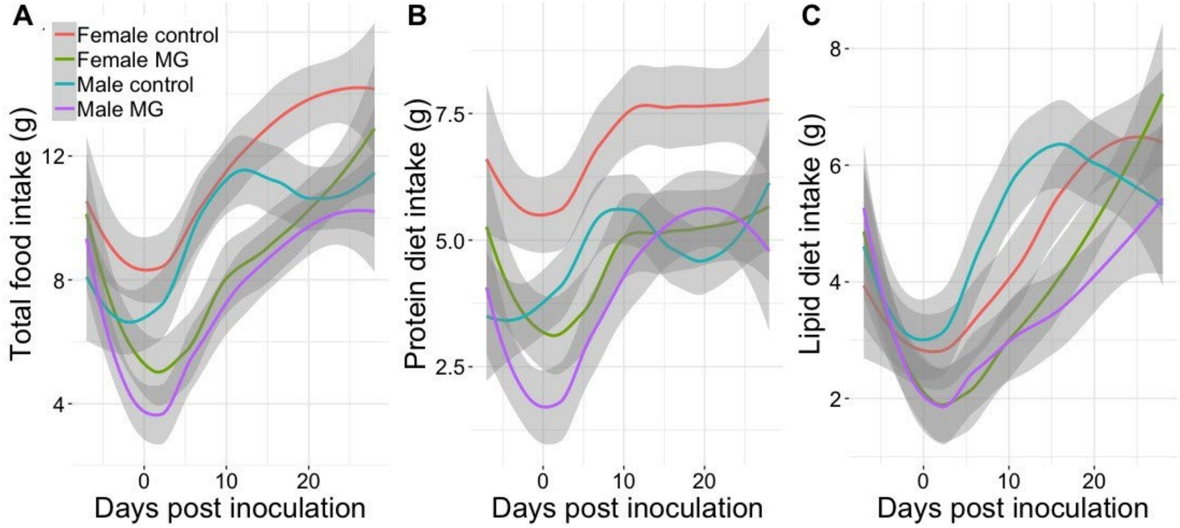


Figure S4 | Differential effect of sex on A) MG-specific antibodies (optical density) and B) log10 transformed MG load over time in only the *Mycoplasma gallisepticum* (MG) exposed canaries (*Serinus canaria domestica*) in experiment two. Points represent raw data with average trend lines surrounded by 95% confidence interval bands in grey.

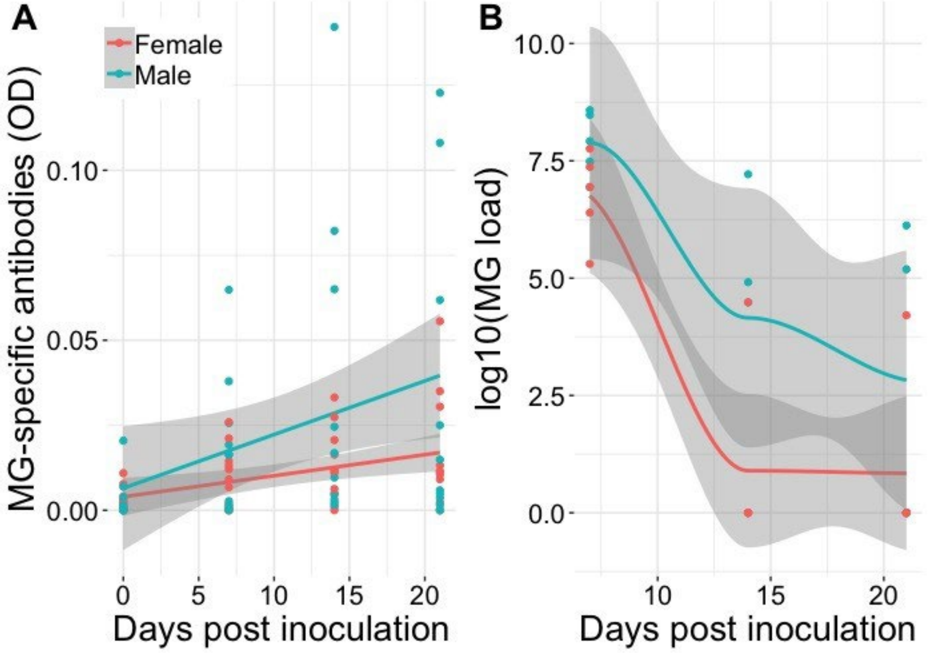


+

Figure S5 | Relative abundance of white blood cells over time from canaries (*Serinus canaria domestica*) in experiment one. Points represent raw data with average trend lines surrounded by 95% confidence interval bands in grey. MG is short for *Mycoplasma gallisepticum.*

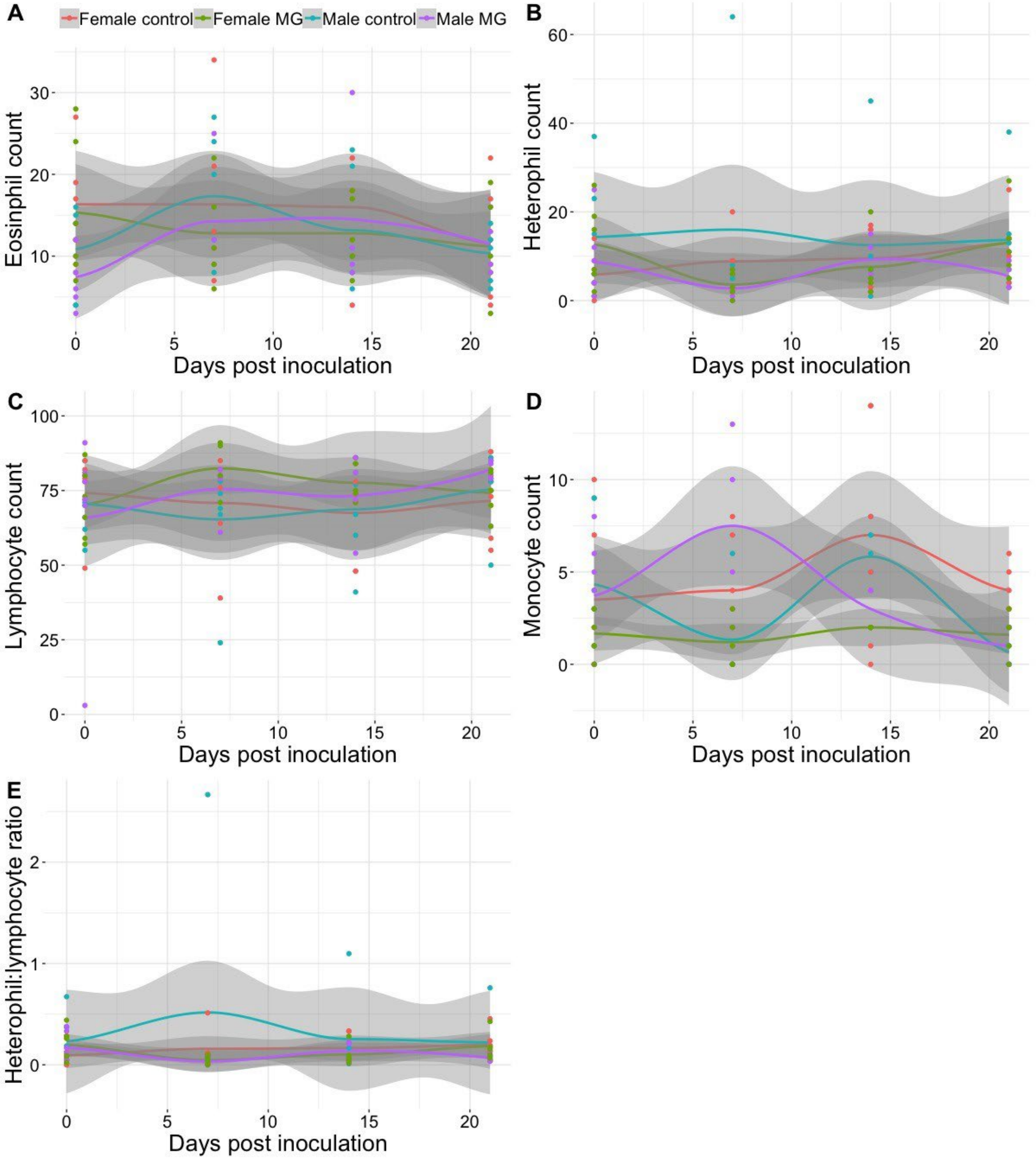

Table S1 | Results from the linear mixed-effects model examining the effect of diet (protein or lipid), *Mycoplasma gallisepticum* (MG) exposure, weeks since MG or sham inoculation, and their interaction on weekly food consumption (g) in canaries (*Serinus canaria domestica*) and ANOVA in experiment one.

| Linear mixed-effects model | | | |
|---|---|---|---|
| | Coefficient | SE | t |
| (Intercept) | 43.012 | 7.234 | 5.946 |
| dietprotein | 7.313 | 9.975 | 0.733 |
| infectionmg | -2.724 | 9.441 | -0.288 |
| week | 0.735 | 1.100 | 0.668 |
| dietprotein:infectionmg | -0.911 | 13.542 | -0.067 |
| dietprotein:week | 0.139 | 1.535 | 0.091 |
| infectionmg:week | -2.316 | 1.538 | -1.506 |
| dietprotein:infectionmg:week | 4.844 | 2.145 | 2.258 |
| ANOVA | | | |
| | Chi squared | df | p value |
| diet | 3.299 | 1 | 0.069 |
| infection | 0.193 | 1 | 0.661 |
| week | 2.800 | 1 | 0.094 |
| diet:infection | 0.399 | 1 | 0.528 |
| diet:week | 5.972 | 1 | 0.015 |
| infection:week | 0.027 | 1 | 0.871 |
| diet:infection:week | 5.099 | 1 | 0.024 |

| Table S2 \| Results from the linear mixed-effects model examining the effect of diet (protein or lipid), *Mycoplasma gallisepticum* (MG) exposure, days since MG or sham inoculation, and their interaction on body mass (g) of canaries (*Serinus canaria domestica*) and ANOVA in experiment one. | | | |
|---|---|---|---|
| Linear mixed-effects model | | | |
| | Coefficient | SE | t |
| (Intercept) | 22.936 | 1.045 | 21.954 |
| Dietprotein | 0.194 | 1.441 | 0.134 |
| infectionMG | -1.717 | 1.410 | -1.217 |
| Day | -0.028 | 0.018 | -1.517 |
| Dietprotein:infectionMG | 0.625 | 1.989 | 0.314 |
| Dietprotein:Day | 0.009 | 0.026 | 0.367 |
| infectionMG:Day | 0.011 | 0.026 | 0.412 |
| Dietprotein:infectionMG:Day | 0.006 | 0.036 | 0.161 |
| ANOVA | | | |
| | Chi squared | df | p value |
| Diet | 0.530 | 1 | 0.466 |
| infection | 1.578 | 1 | 0.209 |
| Day | 3.125 | 1 | 0.077 |
| Diet:infection | 0.137 | 1 | 0.712 |
| Diet:Day | 0.451 | 1 | 0.502 |
| infection:Day | 0.569 | 1 | 0.451 |
| Diet:infection:Day | 0.026 | 1 | 0.872 |

| Table S3 \| Results from the generalized linear mixed-effects model examining the effect of diet (protein or lipid), *Mycoplasma gallisepticum* (MG) exposure, days since MG or sham inoculation, and their interaction on canaries (*Serinus canaria domestica*) hematocrit (%) and ANOVA in experiment one. | | | |
|---|---|---|---|
| Generalized linear mixed-effects model | | | |
| | Coefficient | SE | z |
| (Intercept) | 51.829 | 1.764 | 29.377 |
| DietProtein | 3.100 | 2.497 | 1.241 |
| InfectionMG | 0.123 | 2.294 | 0.054 |
| Day | 0.103 | 0.125 | 0.828 |
| DietProtein:InfectionMG | -0.004 | 3.306 | -0.001 |
| DietProtein:Day | -0.118 | 0.146 | -0.807 |
| InfectionMG:Day | -0.018 | 0.144 | -0.128 |
| DietProtein:InfectionMG:Day | -0.101 | 0.177 | -0.572 |
| ANOVA | | | |
| | Chi squared | df | p value |
| Diet | 1.281 | 1 | 0.258 |
| Infection | 0.156 | 1 | 0.693 |
| Day | 0.062 | 1 | 0.804 |
| Diet:Infection | 0.048 | 1 | 0.826 |
| Diet:Day | 5.310 | 1 | 0.021 |
| Infection:Day | 1.017 | 1 | 0.313 |
| Diet:Infection:Day | 0.327 | 1 | 0.568 |

| Table S4 \| Results from the generalized linear mixed-effects model examining the effect of diet (protein or lipid), *Mycoplasma gallisepticum* (MG) exposure, days since MG or sham inoculation, and their interaction on canaries (*Serinus canaria domestica*) fat score and ANOVA in experiment one. | | | |
|---|---|---|---|
| Generalized linear mixed-effects model | | | |
| | Coefficient | SE | z |
| (Intercept) | 2.087 | 0.291 | 7.182 |
| Dietprotein | -0.246 | 0.401 | -0.615 |
| infectionMG | -0.230 | 0.393 | -0.585 |
| Day | 0.003 | 0.006 | 0.467 |
| Dietprotein:infectionMG | -0.143 | 0.554 | -0.259 |
| Dietprotein:Day | -0.005 | 0.009 | -0.600 |
| infectionMG:Day | -0.003 | 0.009 | -0.283 |
| Dietprotein:infectionMG:Day | -0.003 | 0.013 | -0.204 |
| ANOVA | | | |
| | Chi squared | df | p value |
| Diet | 2.311 | 1 | 0.129 |
| Infection | 1.705 | 1 | 0.192 |
| Day | 0.282 | 1 | 0.595 |
| Diet:Infection | 0.107 | 1 | 0.744 |
| Diet:Day | 1.043 | 1 | 0.307 |
| Infection:Day | 0.373 | 1 | 0.541 |
| Diet:Infection:Day | 0.037 | 1 | 0.848 |

| Table S5 \| Results from the Poisson-distributed generalized additive mixed model of the effect of canary (*Serinus canaria domestica*) diet and the smoothed term days since *Mycoplasma gallisepticum* (MG) exposure on total eye score in experiment one. | | | | |
|---|---|---|---|---|
| Parametric coefficients | | | | |
| | Coefficient | SE | t value | p value |
| (Intercept) | 0.225 | 0.1081 | 2.082 | 0.038 |
| Dietprotein | -0.458 | 0.158 | -2.896 | 0.004 |
| Approximate significance of smooth terms: | | | | |
| | edf | Ref.df | F | p value |
| s(Day) | 1.990 | 1.990 | 130.600 | <0.0001 |

| Table S6 \| Results from the generalized linear model examining the effect of diet (protein or lipid) on recovery time in only *Mycoplasma gallisepticum* (MG) exposed canaries (*Serinus canaria domestica*) and ANOVA in experiment one. | | | |
|---|---|---|---|
| Generalized linear model | | | |
| | Coefficient | SE | t |
| (Intercept) | 25.667 | 3.031 | 8.468 |
| Dietprotein | -8.778 | 4.286 | -2.048 |
| ANOVA | | | |
| | Chi squared | df | p value |
| Diet | 4.194 | 1 | 0.041 |

| Table S7 \| Results from the generalized linear mixed-effects model examining the effect of diet (protein or lipid), days since *Mycoplasma gallisepticum* (MG) exposure, and their interaction on canary (*Serinus canaria domestica*) log10(MG load) and ANOVA in experiment one. | | | |
|---|---|---|---|
| Generalized linear mixed-effects model | | | |
| | Coefficient | SE | z |
| (Intercept) | 8.979 | 0.593 | 15.152 |
| DietProtein | 0.257 | 0.850 | 0.302 |
| day | -0.406 | 0.043 | -9.428 |
| DietProtein:day | 0.010 | 0.061 | 0.158 |
| ANOVA | | | |
| | Chi squared | df | p value |
| Diet | 0.614 | 1 | 0.433 |
| day | 171.051 | 1 | <0.0001 |
| Diet:day | 0.025 | 1 | 0.875 |

| Table S8 \| Results from the generalized linear mixed-effects model examining the effect of diet (protein or lipid), days since *Mycoplasma gallisepticum* (MG) exposure, and their interaction on canary (*Serinus canaria domestica*) MG specific antibodies (optical density) and ANOVA in experiment one. | | | |
|---|---|---|---|
| Generalized linear mixed-effects model | | | |
| | Coefficient | SE | z |
| (Intercept) | 0.004 | 0.014 | 0.307 |
| DietProtein | 0.011 | 0.020 | 0.558 |
| day | 0.004 | 0.001 | 4.289 |
| DietProtein:day | -0.001 | 0.001 | -0.973 |
| ANOVA | | | |
| | Chi squared | df | p value |
| Diet | 0.005 | 1 | 0.945 |
| day | 25.925 | 1 | <0.0001 |
| Diet:day | 0.946 | 1 | 0.331 |

| Table S9 \| Results from the generalized linear mixed-effects model examining the effect of diet (protein or lipid), *Mycoplasma gallisepticum* (MG) exposure, days since MG or sham inoculation, and their interaction on relative canary (*Serinus canaria domestica*) white blood cell counts and ANOVA in experiment one. | | | | | | |
|---|---|---|---|---|---|---|
| | Basophil negative binomial distributed glmm | | | ANOVA | | |
| | Coefficient | SE | z | Chi squared | df | p value |
| (Intercept) | -2.458 | 1.226 | -2.004 | | | |
| DietProtein | 1.872 | 1.284 | 1.457 | 0.582 | 1 | 0.446 |
| InfectionMG | 2.053 | 1.301 | 1.577 | 0.151 | 1 | 0.697 |
| Day | 0.003 | 0.088 | 0.037 | 1.848 | 1 | 0.174 |
| DietProtein:InfectionMG | -3.134 | 1.614 | -1.942 | 0.534 | 1 | 0.465 |
| DietProtein:Day | -0.082 | 0.101 | -0.816 | 0.798 | 1 | 0.372 |
| InfectionMG:Day | -0.206 | 0.129 | -1.596 | 0.345 | 1 | 0.557 |
| DietProtein:InfectionMG:Day | 0.319 | 0.148 | 2.157 | 4.651 | 1 | 0.031 |
| | Eosinophil Poisson distributed glmm | | | ANOVA | | |
| | Coefficient | SE | z | Chi squared | df | p value |
| (Intercept) | 1.400 | 0.303 | 4.615 | | | |
| DietProtein | -0.508 | 0.420 | -1.210 | 4.037 | 1 | 0.045 |
| InfectionMG | -0.069 | 0.405 | -0.171 | 0.038 | 1 | 0.846 |
| Day | 0.037 | 0.016 | 2.333 | 24.635 | 1 | <0.0001 |
| DietProtein:InfectionMG | 0.213 | 0.595 | 0.357 | 0.033 | 1 | 0.856 |
| DietProtein:Day | 0.001 | 0.023 | 0.065 | 0.059 | 1 | 0.808 |
| InfectionMG:Day | 0.006 | 0.021 | 0.281 | 0.006 | 1 | 0.937 |
| DietProtein:InfectionMG:Day | -0.011 | 0.032 | -0.334 | 0.112 | 1 | 0.738 |
| | Heterophil Poisson distributed glmm | | | ANOVA | | |
| | Coefficient | SE | z | Chi squared | df | p value |
| (Intercept) | 2.277 | 0.196 | 11.636 | | | |
| DietProtein | -0.507 | 0.264 | -1.921 | 3.546 | 1 | 0.060 |
| InfectionMG | -0.106 | 0.261 | -0.408 | <0.001 | 1 | 0.998 |
| Day | 0.009 | 0.013 | 0.683 | 8.610 | 1 | 0.003 |
| DietProtein:InfectionMG | 0.252 | 0.386 | 0.652 | 0.137 | 1 | 0.711 |
| DietProtein:Day | 0.019 | 0.017 | 1.099 | 0.980 | 1 | 0.322 |
| InfectionMG:Day | 0.006 | 0.017 | 0.321 | 0.006 | 1 | 0.937 |
| DietProtein:InfectionMG:Day | -0.014 | 0.025 | -0.548 | 0.301 | 1 | 0.583 |
| | Lymphocyte gaussian distributed glmm | | | ANOVA | | |

| | Coefficient | SE | z | Chi squared | df | p value |
|---|---|---|---|---|---|---|
| (Intercept) | 84.480 | 2.456 | 34.393 | | | |
| DietProtein | 4.069 | 3.186 | 1.277 | 9.801 | 1 | 0.002 |
| InfectionMG | -1.318 | 3.266 | -0.404 | 0.034 | 1 | 0.854 |
| Day | -0.593 | 0.180 | -3.295 | 24.800 | 1 | <0.0001 |
| DietProtein:InfectionMG | -1.239 | 4.558 | -0.272 | 0.812 | 1 | 0.368 |
| DietProtein:Day | 0.191 | 0.233 | 0.818 | 0.529 | 1 | 0.467 |
| InfectionMG:Day | 0.235 | 0.240 | 0.979 | 0.934 | 1 | 0.334 |
| DietProtein:InfectionMG:Day | -0.142 | 0.335 | -0.426 | 0.181 | 1 | 0.670 |
| | Monocyte Poisson distributed glmm | | | ANOVA | | |
| | Coefficient | SE | z | Chi squared | df | p value |
| (Intercept) | 0.630 | 0.419 | 1.503 | | | |
| DietProtein | 0.074 | 0.526 | 0.140 | 0.365 | 1 | 0.546 |
| InfectionMG | 0.084 | 0.543 | 0.155 | 0.805 | 1 | 0.370 |
| Day | 0.040 | 0.025 | 1.597 | 3.143 | 1 | 0.076 |
| DietProtein:InfectionMG | 0.214 | 0.728 | 0.293 | 0.426 | 1 | 0.514 |
| DietProtein:Day | -0.006 | 0.032 | -0.187 | 0.002 | 1 | 0.968 |
| InfectionMG:Day | -0.042 | 0.034 | -1.237 | 2.621 | 1 | 0.105 |
| DietProtein:InfectionMG:Day | 0.010 | 0.045 | 0.225 | 0.051 | 1 | 0.822 |
| | Heterophil:lymphocyte ratio binomial distributed glmm | | | ANOVA | | |
| | Coefficient | SE | z | Chi squared | df | p value |
| (Intercept) | -2.266 | 0.188 | -12.086 | | | |
| DietProtein | -0.547 | 0.256 | -2.140 | 4.951 | 1 | 0.026 |
| InfectionMG | -0.034 | 0.254 | -0.134 | 0.025 | 1 | 0.873 |
| Day | 0.025 | 0.008 | 3.196 | 40.620 | 1 | <0.0001 |
| DietProtein:InfectionMG | 0.322 | 0.367 | 0.877 | 0.476 | 1 | 0.490 |
| DietProtein:Day | 0.007 | 0.011 | 0.653 | 0.124 | 1 | 0.724 |
| InfectionMG:Day | -0.004 | 0.010 | -0.421 | 1.222 | 1 | 0.269 |
| DietProtein:InfectionMG:Day | -0.009 | 0.015 | -0.573 | 0.328 | 1 | 0.567 |

| Table S10 \| Results from the linear mixed-effects model examining the effect of sex, *Mycoplasma gallisepticum* (MG) exposure, days since MG or sham inoculation, and their interaction on canary (*Serinus canaria domestica*) weekly total food consumption (g) and ANOVA in experiment two. | | | |
|---|---|---|---|
| Linear mixed-effects model | | | |
| | Coefficient | SE | t |
| (Intercept) | 59.944 | 7.298 | 8.214 |
| sexM | -7.322 | 10.320 | -0.709 |
| infectionmg | -17.122 | 10.412 | -1.644 |
| Week | 10.165 | 1.867 | 5.445 |
| sexM:infectionmg | 0.291 | 14.795 | 0.020 |
| sexM:Week | -3.225 | 2.640 | -1.222 |
| infectionmg:Week | -1.406 | 2.742 | -0.513 |
| sexM:infectionmg:Week | 3.373 | 3.980 | 0.847 |
| ANOVA | | | |
| | Chi squared | df | p value |
| sex | 2.643 | 1 | 0.104 |
| infection | 6.657 | 1 | 0.010 |
| Week | 77.367 | 1 | <0.0001 |
| sex:infection | 0.245 | 1 | 0.621 |
| sex:Week | 0.777 | 1 | 0.378 |
| infection:Week | 0.010 | 1 | 0.922 |
| sex:infection:Week | 0.718 | 1 | 0.397 |

| Table S11 \| Results from the linear mixed-effects model examining the effect of sex, Mycoplasma gallisepticum (MG) exposure, days since MG or sham innoculation, and their interaction on canary (Serinus canaria domestica) weekly protein diet consumption (g) and ANOVA in experiment two. | | | |
|---|---|---|---|
| Linear mixed-effects model | | | |
| | Coefficient | SE | t |
| (Intercept) | -2.266 | 0.188 | -12.086 |
| sexM | -0.547 | 0.256 | -2.140 |
| infectionmg | -0.034 | 0.254 | -0.134 |
| Week | 0.025 | 0.008 | 3.196 |
| sexM:infectionmg | 0.322 | 0.367 | 0.877 |
| sexM:Week | 0.007 | 0.011 | 0.653 |
| infectionmg:Week | -0.004 | 0.010 | -0.421 |
| sexM:infectionmg:Week | -0.009 | 0.015 | -0.573 |
| ANOVA | | | |
| | Chi squared | df | p value |
| sex | 5.769 | 1 | 0.016 |
| infection | 4.237 | 1 | 0.040 |
| Week | 26.824 | 1 | <0.0001 |
| sex:infectionmg | 1.127 | 1 | 0.288 |
| sex:Week | 0.044 | 1 | 0.834 |
| infection:Week | 0.842 | 1 | 0.359 |
| sex:infection:Week | 1.492 | 1 | 0.222 |

| Table S12 \| Results from the linear mixed-effects model examining the effect of sex, Mycoplasma gallisepticum (MG) exposure, days since MG or sham innoculation, and their interaction on canary (Serinus canaria domestica) weekly lipid diet consumption (g) and ANOVA in experiment two. | | | |
|---|---|---|---|
| Linear mixed-effects model | | | |
| | Coefficient | SE | t |
| (Intercept) | 19.094 | 4.300 | 4.440 |
| sexM | 6.071 | 6.081 | 0.998 |
| infectionmg | -3.276 | 6.148 | -0.533 |
| Week | 6.475 | 1.260 | 5.139 |
| sexM:infectionmg | -3.539 | 8.747 | -0.405 |
| sexM:Week | -1.995 | 1.782 | -1.120 |
| infectionmg:Week | -0.997 | 1.846 | -0.540 |
| sexM:infectionmg:Week | -0.201 | 2.675 | -0.075 |
| ANOVA | | | |
| | Chi squared | df | p value |
| sex | 0.023 | 1 | 0.879 |
| infection | 3.751 | 1 | 0.053 |
| Week | 57.503 | 1 | <0.0001 |
| sex:infection | 0.292 | 1 | 0.589 |
| sex:Week | 2.461 | 1 | 0.117 |
| infection:Week | 0.669 | 1 | 0.413 |
| sex:infection:Week | 0.006 | 1 | 0.940 |

| Table S13 \| Results from the linear mixed-effects model examining the effect of sex, *Mycoplasma gallisepticum* (MG) exposure, days since MG or sham inoculation, and their interaction on canary (*Serinus canaria domestica*) mass (g) and ANOVA in experiment two. | | | |
|---|---|---|---|
| Linear mixed-effects model | | | |
| | Coefficient | SE | t |
| (Intercept) | 25.667 | 1.657 | 15.493 |
| sexM | -5.091 | 2.343 | -2.173 |
| infectionmg | -4.566 | 2.350 | -1.943 |
| day | -0.021 | 0.025 | -0.815 |
| sexM:infectionmg | 3.570 | 3.268 | 1.092 |
| sexM:day | 0.038 | 0.036 | 1.067 |
| infectionmg:day | -0.008 | 0.037 | -0.213 |
| sexM:infectionmg:day | -0.060 | 0.054 | -1.115 |
| ANOVA | | | |
| | Chi squared | df | p value |
| sex | 3.812 | 1 | 0.051 |
| infection | 4.040 | 1 | 0.044 |
| day | 1.716 | 1 | 0.190 |
| sex:infection | 0.760 | 1 | 0.383 |
| sex:day | 0.197 | 1 | 0.657 |
| infection:day | 1.811 | 1 | 0.178 |
| sex:infection:day | 1.243 | 1 | 0.265 |

Table S14 | Results from the generalized linear mixed-effects model examining the effect of sex, *Mycoplasma gallisepticum* (MG) exposure, days since MG or sham inoculation, and their interaction on canary (*Serinus canaria domestica*) hematocrit (%) and ANOVA in experiment two.

| Generalized linear mixed-effects model | | | |
|---|---|---|---|
| | Coefficient | SE | z |
| (Intercept) | 54.850 | 2.157 | 25.427 |
| sexM | 8.783 | 3.051 | 2.879 |
| infectionmg | -0.851 | 3.083 | -0.276 |
| day | 0.014 | 0.094 | 0.152 |
| sexM:infectionmg | -4.839 | 4.286 | -1.129 |
| sexM:day | 0.040 | 0.133 | 0.305 |
| infectionmg:day | -0.006 | 0.143 | -0.040 |
| sexM:infectionmg:day | -0.152 | 0.204 | -0.746 |
| ANOVA | | | |
| | Chi squared | df | p value |
| sex | 10.255 | 1 | 0.001 |
| infection | 4.247 | 1 | 0.039 |
| day | 0.0001 | 1 | 0.992 |
| sex:infection | 2.546 | 1 | 0.111 |
| sex:day | 0.056 | 1 | 0.812 |
| infection:day | 0.625 | 1 | 0.429 |
| sex:infection:day | 0.557 | 1 | 0.455 |

| Table S15 \| Results from the linear mixed-effects model examining the effect of sex, *Mycoplasma gallisepticum* (MG) exposure, days since MG or sham inoculation, and their interaction on canary (*Serinus canaria domestica*) fat score and ANOVA in experiment two. | | | |
|---|---|---|---|
| Linear mixed-effects model | | | |
| | Coefficient | SE | t |
| (Intercept) | 2.025 | 0.286 | 7.087 |
| sexM | -1.097 | 0.404 | -2.715 |
| infectionmg | -0.319 | 0.407 | -0.786 |
| day | -0.002 | 0.006 | -0.256 |
| sexM:infectionmg | 0.338 | 0.566 | 0.598 |
| sexM:day | 0.023 | 0.009 | 2.532 |
| infectionmg:day | -0.009 | 0.009 | -0.921 |
| sexM:infectionmg:day | -0.010 | 0.014 | -0.720 |
| ANOVA | | | |
| | Chi squared | df | p value |
| sex | 6.426 | 1 | 0.011 |
| infection | 1.429 | 1 | 0.232 |
| day | 1.121 | 1 | 0.290 |
| sex:infection | 0.152 | 1 | 0.696 |
| sex:day | 7.483 | 1 | 0.006 |
| infection:day | 3.805 | 1 | 0.051 |
| sex:infection:day | 0.518 | 1 | 0.472 |

| Table S16 \| Results from the Poisson-distributed generalized additive mixed model of the effect of canary (*Serinus canaria domestica*) diet and the smoothed term days since *Mycoplasma gallisepticum* (MG) exposure on total eye score in experiment two. | | | | |
|---|---|---|---|---|
| Parametric coefficients | | | | |
| | Coefficient | SE | t value | p value |
| (Intercept) | -1.2417 | 0.1794 | -6.92 | <0.0001 |
| SexM | 1.9289 | 0.221 | 8.727 | <0.0001 |
| Approximate significance of smooth terms | | | | |
| | edf | Ref.df | F | p-value |
| s(Day) | 1.836 | 1.836 | 3.964 | 0.068 |

| Table S17 \| Results from the generalized linear mixed-effects model examining the effect of sex, days since *Mycoplasma gallisepticum* (MG) exposure, and their interaction on canary (*Serinus canaria domestica*) log10(MG load) and ANOVA in experiment two. | | | |
|---|---|---|---|
| Generalized linear mixed-effects model | | | |
| | Coefficient | SE | z |
| (Intercept) | 8.744 | 1.322 | 6.612 |
| sexm | 1.347 | 1.893 | 0.712 |
| day | -0.422 | 0.080 | -5.260 |
| sexm:day | 0.061 | 0.118 | 0.514 |
| ANOVA | | | |
| | Chi squared | df | p value |
| sex | 4.593 | 1 | 0.032 |
| day | 44.864 | 1 | <0.0001 |
| sex:day | 0.264 | 1 | 0.608 |

| Table S18 \| Results from the generalized linear mixed-effects model examining the effect of sex, days since *Mycoplasma gallisepticum* (MG) exposure, and their interaction on canary (*Serinus canaria domestica*) MG specific antibody level (optical density) and ANOVA in experiment two. | | | |
|---|---|---|---|
| Generalized linear mixed-effects model | | | |
| | Coefficient | SE | z |
| (Intercept) | 0.004 | 0.007 | 0.596 |
| sexm | 0.003 | 0.010 | 0.263 |
| day | 0.001 | 0.001 | 1.770 |
| sexm:day | 0.001 | 0.001 | 1.888 |
| ANOVA | | | |
| | Chi squared | df | p value |
| sex | 2.466 | 1 | 0.116 |
| day | 18.020 | 1 | <0.0001 |
| sex:day | 3.563 | 1 | 0.059 |

| Table S19 \| Results from the generalized linear mixed-effects model examining the effect of diet (protein or lipid), *Mycoplasma gallisepticum* (MG) exposure, days since MG or sham inoculation, and their interaction on relative canary (*Serinus canaria domestica*) white blood cell counts and ANOVA in experiment two. | | | | | | |
|---|---|---|---|---|---|---|
| | Eosinophil gaussian distributed glmm | | | ANOVA | | |
| | Coefficient | SE | z | Chi squared | df | p value |
| (Intercept) | 13.767 | 2.368 | 5.813 | | | |
| SexM | 3.500 | 3.349 | 1.045 | 1.601 | 1 | 0.206 |
| InfectionMG | -4.845 | 3.282 | -1.476 | 1.333 | 1 | 0.248 |
| Day | -0.081 | 0.120 | -0.674 | 1.987 | 1 | 0.159 |
| SexM:InfectionMG | 2.503 | 4.705 | 0.532 | 0.009 | 1 | 0.925 |
| SexM:Day | -0.131 | 0.170 | -0.771 | 3.404 | 1 | 0.065 |
| InfectionMG:Day | 0.258 | 0.184 | 1.402 | 1.171 | 1 | 0.279 |
| SexM:InfectionMG:Day | -0.230 | 0.255 | -0.905 | 0.818 | 1 | 0.366 |
| | Heterophil Poisson distributed glmm | | | ANOVA | | |
| | Coefficient | SE | z | Chi squared | df | p value |
| (Intercept) | 1.730 | 0.321 | 5.383 | | | |
| SexM | 0.645 | 0.430 | 1.501 | 0.010 | 1 | 0.921 |
| InfectionMG | 0.232 | 0.462 | 0.504 | 0.604 | 1 | 0.437 |
| Day | 0.035 | 0.015 | 2.399 | 2.763 | 1 | 0.096 |

| | | | | | | |
|---|---|---|---|---|---|---|
| SexM:InfectionMG | -0.748 | 0.628 | -1.191 | 1.342 | 1 | 0.247 |
| SexM:Day | -0.037 | 0.019 | -1.941 | 4.409 | 1 | 0.036 |
| InfectionMG:Day | -0.013 | 0.023 | -0.538 | 0.147 | 1 | 0.702 |
| SexM:InfectionMG:Day | 0.012 | 0.033 | 0.377 | 0.142 | 1 | 0.706 |
| | Lymphocyte gaussian distributed glmm | | | ANOVA | | |
| | Coefficient | SE | z | Chi squared | df | p value |
| (Intercept) | 72.700 | 6.623 | 10.977 | | | |
| SexM | -5.417 | 9.367 | -0.578 | 0.449 | 1 | 0.503 |
| InfectionMG | 1.198 | 9.390 | 0.128 | 0.002 | 1 | 0.969 |
| Day | -0.162 | 0.194 | -0.834 | 0.592 | 1 | 0.441 |
| SexM:InfectionMG | -3.499 | 13.062 | -0.268 | 0.174 | 1 | 0.677 |
| SexM:Day | 0.417 | 0.275 | 1.517 | 2.567 | 1 | 0.109 |
| InfectionMG:Day | 0.187 | 0.287 | 0.652 | 0.206 | 1 | 0.650 |
| SexM:InfectionMG:Day | -0.197 | 0.418 | -0.470 | 0.221 | 1 | 0.638 |
| | Monocyte Poisson distributed glmm | | | ANOVA | | |
| | Coefficient | SE | z | Chi squared | df | p value |
| (Intercept) | 1.233 | 0.298 | 4.133 | | | |
| SexM | 0.028 | 0.434 | 0.064 | 0.008 | 1 | 0.927 |
| InfectionMG | -0.619 | 0.484 | -1.279 | 1.146 | 1 | 0.284 |
| Day | 0.019 | 0.019 | 1.039 | 0.522 | 1 | 0.470 |
| SexM:InfectionMG | 0.908 | 0.635 | 1.431 | 4.372 | 1 | 0.037 |
| SexM:Day | -0.052 | 0.030 | -1.716 | 4.516 | 1 | 0.034 |
| InfectionMG:Day | -0.013 | 0.032 | -0.392 | 0.151 | 1 | 0.698 |
| SexM:InfectionMG:Day | 0.008 | 0.046 | 0.167 | 0.028 | 1 | 0.868 |
| | Heterophil:lymphocyte ratio binomial distributed glmm | | | ANOVA | | |
| | Coefficient | SE | z | Chi squared | df | p value |
| (Intercept) | -2.570 | 0.345 | -7.448 | | | |
| SexM | 0.702 | 0.481 | 1.459 | 0.0002 | 1 | 0.988 |
| InfectionMG | 0.329 | 0.485 | 0.678 | 0.104 | 1 | 0.747 |
| Day | 0.036 | 0.009 | 3.829 | 4.926 | 1 | 0.026 |
| SexM:InfectionMG | -0.714 | 0.680 | -1.050 | 0.682 | 1 | 0.409 |
| SexM:Day | -0.044 | 0.013 | -3.480 | 13.605 | 1 | 0.0002 |
| InfectionMG:Day | -0.016 | 0.014 | -1.154 | 0.534 | 1 | 0.465 |
| SexM:InfectionMG:Day | 0.019 | 0.021 | 0.910 | 0.829 | 1 | 0.363 |
